# A targeted bispecific TGFBR2 antagonist antibody demonstrates cell selectivity and enhanced potency on human fibroblasts

**DOI:** 10.64898/2026.08.11.744117

**Authors:** Russell B. Fletcher, Hui Chen, Yorick Post, Yiran Yang, Navrose Dhaliwal, Yongfeng Fan, Trevor Fisher, Sungjin Lee, Nicholas Suen, Molly Smith, Nick Downs, Jay Ye, Jonathan Karr, Sarah G. Hymowitz, Mridula K. Ray, Chenggang Lu, Yang Li

## Abstract

Beyond its critical roles in development and tissue homeostasis, TGFβ signaling promotes key aspects of cancer progression and is a primary driver of fibrosis. Although blocking TGFβ signaling has great therapeutic potential for cancer and fibrotic diseases such as idiopathic pulmonary fibrosis (IPF), preclinical and clinical studies revealed that non-specific alteration of the pathway can have severe adverse consequences; therefore, inhibiting TGFβ signaling in a cell-type specific manner may avoid systemic toxic effects while preserving potential therapeutic effects. The parasitic helminth *Heligmosomoides polygyrus* has evolved cell-type-targeted modulators of TGFβ signaling. With insights from the development of other targeted signaling modulators and using the worm proteins as a guide, we sought to develop a human-fibroblast-targeted TGFBR2 antagonist. Here, we report mechanistic insights into the targeted worm TGFBR2 antagonist TGM6 and fusion proteins containing the TGM6 targeting domains. We created a bispecific antibody TGFBR2 antagonist that binds PDGFRA as a targeting receptor and demonstrates cell selectivity and enhanced potency in fibroblasts. Our findings suggest a viable path for developing targeted TGFβ signaling antagonists as therapeutics for cancer and tissue fibrosis.

## Introduction

The TGFβ cytokine family has important pleiotropic functions in embryonic development and tissue homeostasis (Massague & Sheppard, 2023). Although TGFβ signaling is an attractive therapeutic target for cancer and fibrosis, attempts to modulate it for therapeutic benefit have been fraught with difficulty. Beyond promoting metastases, immune system evasion, and immunosuppression in cancer, the pathway promotes fibrosis across a range of tissues by inducing myofibroblast states and extracellular matrix (ECM) deposition, thus severely compromising tissue function and limiting tumor accessibility to therapeutics (Derynck et al., 2021; Frangogiannis, 2020; Massague, 2008; Massague & Sheppard, 2023). Idiopathic pulmonary fibrosis (IPF) is an especially grievous example of a fibrotic lung disease where it is thought that aberrant epithelial cells are involved in its initiation, leading to activation of ECM secreting myofibroblasts, and TGFβ signaling is a central driver (Fernandez & Eickelberg, 2012; Pardo & Selman, 2016; Parimon et al., 2020; Wolters et al., 2018). Currently approved drugs for IPF (Nintedanib and Pirfenidone) only weakly limit functional decline and have undesirable side-effects; only lung transplantation has been shown to increase patient survival (Anderson et al., 2024; George et al., 2019; Riddell et al., 2020; Toth et al., 2025). Therefore, there is a need for therapies that directly impact fibrogenic drivers of the disease.

Blocking TGFβ signaling has promise for treating IPF, but untargeted TGFβ signaling inhibition has detrimental consequences, as illustrated in attempts to develop antagonist/inhibitor therapeutic candidates. In preclinical studies, animals treated with small molecule inhibitors of TGFBR1 or an anti-TGFB1,2,3 antibody displayed cardiovascular damage such as hemorrhaging and cardiac valve degeneration (Anderton et al., 2011; Herbertz et al., 2015; Mitra et al., 2020). Moreover, TGFBR2 is an obligate and specific receptor for the three TGFB ligands, and in a clinical trial of patients with solid tumors, treatment with a high affinity TGFBR2 blocking antibody led to an exacerbated immune response that halted development of the drug (Tolcher et al., 2017). More recently, the BEACON-IPF Phase 2b/3 trial of bexotegrast (Wuyts et al., 2026), a small molecule inhibitor of αvβ8 and αvβ1 integrins that blocks TGFβ ligand activation, was halted due to safety concerns over IPF-related adverse events (Pliant Therapeutics, 2025), suggesting that more precise targeting of the TGFβ signaling blockade might be necessary to avoid acute exacerbations in patients with IPF. In line with this, a TGFB3-specific blocking antibody attenuated fibrosis in a preclinical model of pulmonary fibrosis with minimal toxicity (Sun et al., 2024), although it may not be as effective as a TGFBR2 blockade due to the potential activity of other TGFB ligands.

Limiting blockade of TGFβ signaling to specific cell types and avoiding cells associated with adverse consequences may enable therapeutic success in targeting this powerful pathway. Several examples exist that couple a cell-type-specific antigen binder to target a cell signaling modulator. For example, a Phase 1 trial has recently tested a hepatocyte-targeted enhancer of Wnt signaling that uses ASGR1 binding to localize R-SPONDIN (RSPO) activity to hepatocytes, redirecting RSPO to ASGR1+ cells instead of LGR+ cells (Sampathkumar et al., 2024; Zhang et al., 2020). In a related example, a WNT ligand surrogate that uses two inactive halves, each targeting separate epitopes on a hepatocyte-specific receptor, has been targeted to hepatocytes for subsequent activation (Chen et al., 2023). Similarly, Bicara therapeutics recently reported Phase 1b results for Ficerafusp Alfa (BCA101), an EGFR targeted TGFB1,3 trap (Boreddy et al., 2023), that in combination with the PD-1 (PDCD1) blocking antibody pembrolizumab was safe and appeared to have beneficial effects on HNSCC tumors (Ferrarotto R, 2025). Nature has also provided several elegant solutions: the parasitic helminth *Heligmosomoides polygyrus* resides in the mouse gut and has evolved targeted agonists and antagonists of TGFβ signaling which manipulate its local environment and the host response: these molecules, termed TGFB mimetics (TGMs) contain modular domains that bind TGFBR1 and/or TGFBR2 and other cell surface receptors that lead to cell selective pathway activation or antagonism (Johnston et al., 2017; Singh et al., 2025; Smyth et al., 2018; White et al., 2025). TGM6 is a TGFBR2 antagonist coupled to a cell-type targeting domain (Smyth et al., 2018; White et al., 2025).

Mindful of the risk of systemic, non-specific TGFβ signaling blockade, we set out to develop an antibody-based fibroblast-targeted TGFBR2 antagonist. First, we investigated the activity of the worm TGFBR2 antagonist (TGM6) and explored fusion proteins comprising an antibody fragment that binds TGFBR2 coupled to the TGM6 targeting domains. We then designed a bi-specific antibody-based antagonist that binds TGFBR2 and PDGFRA, a receptor expressed on pulmonary fibroblasts and thought to be critical in pulmonary fibrosis development (Tsukui et al., 2024). The PDGFRA-targeted TGFBR2 antagonist displayed cell selectivity and enhanced antagonist activity, with ∼300-fold greater potency relative to an untargeted competitive antagonist in human fetal pulmonary fibroblasts. This *in vitro* proof-of-concept demonstrated that a cell-type-targeted TGFBR2 antagonist can achieve cell type specificity and enhanced potency, thereby supporting a potential path to developing therapeutics that selectively inhibit TGFβ signaling.

## Results & Discussion

### TGM6 characterization

The parasitic helminth *Heligmosomoides polygyrus* evolved a family of TGM proteins that either activate or inhibit TGFβ signaling (Maizels & Newfeld, 2023; Singh et al., 2025; Smyth et al., 2018; White et al., 2025). TGM6 is a TGFBR2 antagonist that demonstrates selective activity in mouse fibroblasts while sparing immune cells. TGM6 is comprised of three modular domains, the TGFBR2 binding domain D3 as well as D4 and D5 (D4D5) which are thought to enable cell selective activity by binding one or more cell-specific co-receptors (White et al., 2025). How TGM6 achieves highly potent sub-nanomolar inhibitory activity on fibroblasts with a modest affinity toward TGFBR2 (reported K_D_ of 220-440 nM for TGM6-D3 binding to TGFBR2) was not previously elucidated (White et al., 2025). To understand the mechanism of TGM6 action as a prelude to building a selective antagonist for human fibroblasts, we expressed and characterized recombinant TGM6. This TGM6 protein was purified to a single species (Figure 1A). Biolayer interferometry indicated that it bound both mouse and human TGFBR2 with ∼90-fold higher affinity for the mouse receptor (K_D_ = 1.4 nM versus 122 nM) (Figure 1B). Furthermore, in agreement with its initial characterization by White et al. (2025), TGM6 competed with a TGFB for binding to TGFBR2 (Figure 1C).

**Figure 1.**
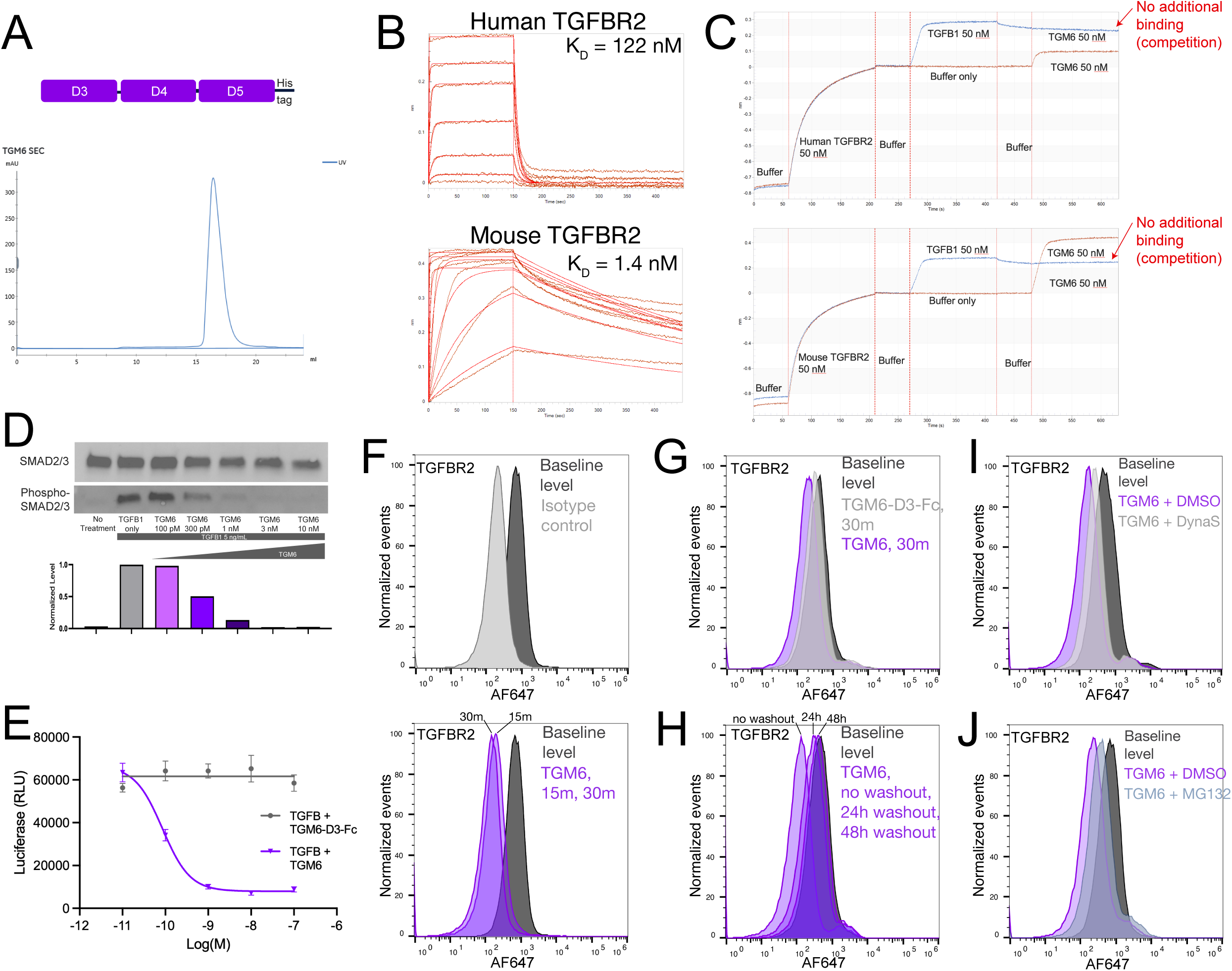
TGM6 binding and activity. (A) The three-module domain layout and His-tag position of TGM6. The protein was a single species according to SEC. (B) Biolayer interferometry (BLI)-based assays show TGM6 binding to mouse and human TGFBR2 with the indicated affinity K_D_ values. (C) TGM6 competes with TGFB1 for binding to mouse and human TGFBR2 by BLI. (D, E) TGM6 inhibits TGFB1-induced signaling, as indicated by Western blotting for phospho-SMAD2/3 (D) and by a TGFB SMAD-binding element (SBE) luciferase reporter (E). (F-I) TGFBR2 detected by flow cytometry in NIH-3T3 cells for normal, untreated levels (F, upper), and 15 or 30 minutes after TGM6 treatment (F, lower), after 30 minutes of TGM6-D3-Fc or TGM6 treatment (G), or after 30 minutes of TGM6 treatment followed by the indicated washout period after TGM6 removal (H), and after a 30-minute TGM6 treatment in the presence of the dynamin inhibitor Dynasore (I) or the proteosome inhibitor MG132 (J). All functional assays were repeated at least three times.

Functional *in vitro* assays indicated that TGM6 antagonized TGFB1-induced signaling in mouse fibroblasts with an IC50 around 300 pM, whereas the TGM6-D3 domain alone had no inhibitory activity (Figure 1D, E), confirming that TGM6 antagonist activity required the TGM6-D4D5 domains. We considered two potential explanations for the D4D5-dependent activity of TGM6. We reasoned that coupling a TGFBR2 binder with a binder to a cell-type-specific targeting receptor might increase the potency of the molecule via cooperativity. In a second, non-mutually exclusive model, TGM6 might act via a lysosomal degradation mechanism similar to what some have referred to as LYTAC (Lysosome-targeting chimera) (Banik et al., 2020) in which target receptor engagement leads to rapid internalization and degradation of the TGFBR2 receptor.

We investigated surface levels of TGFBR2 in the mouse NIH-3T3 fibroblast line through flow cytometry. NIH-3T3 cells express ample TGFBR2 on the cell surface, and these levels are quickly decreased to isotype control levels within 15 minutes of TGM6 treatment (Figure 1F). In contrast, the TGM6-D3 domain alone causes only a minor decrease in TGFBR2 levels after a 30-minute treatment (Figure 1G). Next, we tested the time required for surface TGFBR2 levels to replenish after a 30-minute TGM6 treatment. No notable restoration was observed after 2- or 4-hours (data not shown), but the surface receptor levels returned to approximately 2/3 the initial levels after a 24-hour washout and nearly completely recovered after 48 hours (Figure 1H). The robust, rapid decrease in TGFBR2 levels along with the delayed replenishment suggested that TGM6 might act via a LYTAC-like mechanism. In agreement with this possibility, both a dynamin inhibitor and proteosome inhibitor partially blocked the TGM6-mediated removal of TGFBR2 from the cell surface (Figure 1I, J).

### Bispecific fusion protein TGFBR2 antagonist

While TGM6 showed intriguing proof-of-concept for this approach to modulating TGFβ signaling, it is unlikely to be a successful therapeutic due to the high potential for immunogenicity from helminth-derived protein sequences. Thus, we explored whether we could generate a more drug-like antibody-based molecule targeting human fibroblasts. First, we assessed whether the TGM6 D3 domain could be replaced by a TGFBR2 binding antibody fragment. We selected a variable heavy chain (VH) binder reported to bind TGFBR2 (clone VH Dom23h-271-12, WO2012/093125A1) and constructed a bi-specific fusion molecule by combining VH 271-12 with the TGM6-D4D5 domain (Figure 2A). The VH 271-12 had an affinity for mouse TGFBR2 approximately 20-fold lower than that of the TGM6-D3 domain; moreover, like the TGM6-D3 domain, it competed with TGFB ligand for TGFBR2 binding (Figure 2 B, C). In mouse NIH-3T3 fibroblasts, the VH 271-12 + TGM6-D4D5 bispecific fusion molecule displayed antagonist activity comparable to that of TGM6 (Figure 2D), and it removed TGFBR2 from the cell surface in a similar manner (Figure 2E). While TGM6 has no to very limited activity in human cells, the VH 271-12 + TGM6-D4D5 fusion protein blocked TGFB-induced signaling in human fibroblasts (Figure 2F). These results not only further supported a LYTAC-like mechanism but also indicated that the affinity range for effective TGFBR2 binders can be significantly weaker than the TGM6-D3 domain.

**Figure 2.**
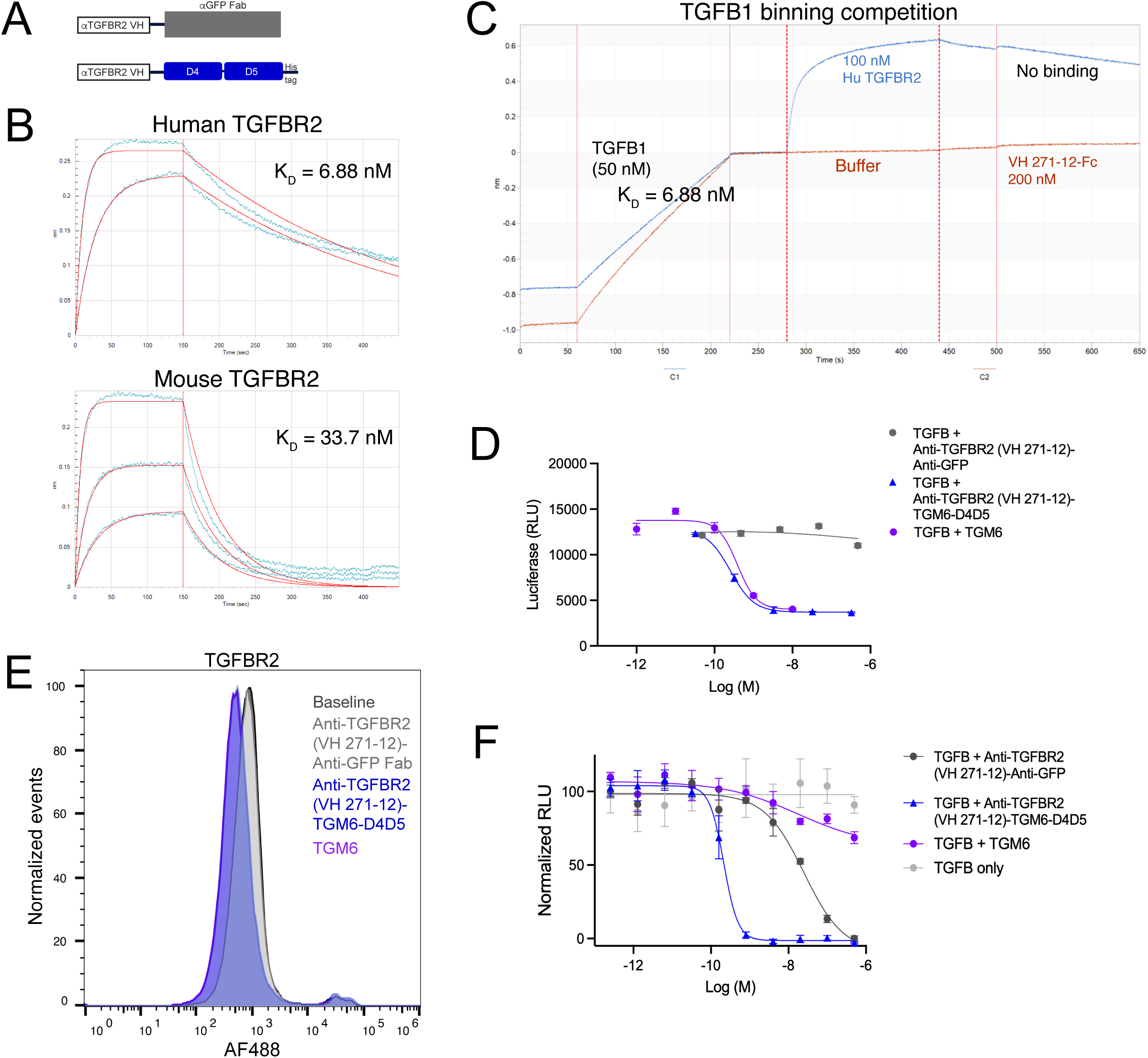
Fusion proteins with the TGM6-D4D5 targeting domains have robust activity. (A) Design of the control and TGM6-D4D5 fusion proteins. (B, C) The binder VH 271-12 binds human and mouse TGFBR2 with the indicated affinity K_D_ values and competes with TGFB1 for TGFBR2 binding as assessed by BLI. (D) Activity of the VH 271-12-TGM6-D4D5 fusion protein in the SBE luciferase reporter assay in NIH-3T3 cells. (E) Surface levels of TGFBR2 measured by flow cytometry after fusion protein treatments in NIH-3T3 cells. (F) Activity of the VH 271-12-TGM6-D4D5 fusion protein in the SBE luciferase reporter assay in human MRC5 fibroblasts. The activity assay was repeated in mouse cells in triplicate and in human cells in duplicate.

### In vitro proof-of-concept of an antibody based human fibroblast selective TGFBR2 antagonist

Next, we replaced the TGM6-D4D5 domains with an antibody binder to generate a fully antibody-based molecule with reduced potential for immunogenicity while enhancing functional selectivity for human fibroblasts. Although TGM6 has robust activity in mouse fibroblasts, it does not work in human cells, and the identity of the receptor(s) bound by the TGM6-D4D5 domains was unknown at the time of this study (White et al., 2025). PDGFRA is a signaling receptor enriched in fibroblasts, and PDGFRA-positive pulmonary fibroblasts have recently been demonstrated to play a crucial role in pulmonary fibrosis (Tsukui et al., 2024), making it a favorable candidate targeting receptor with relevance to IPF. For an untargeted control, we combined the anti-TGFBR2 (271-12) VH with an anti-GFP Fab and constructed a targeted bispecific antagonist by combining VH 271-12 with an anti-PDGFRA Fab (US2014/8697664) (Figure 3A).

**Figure 3.**
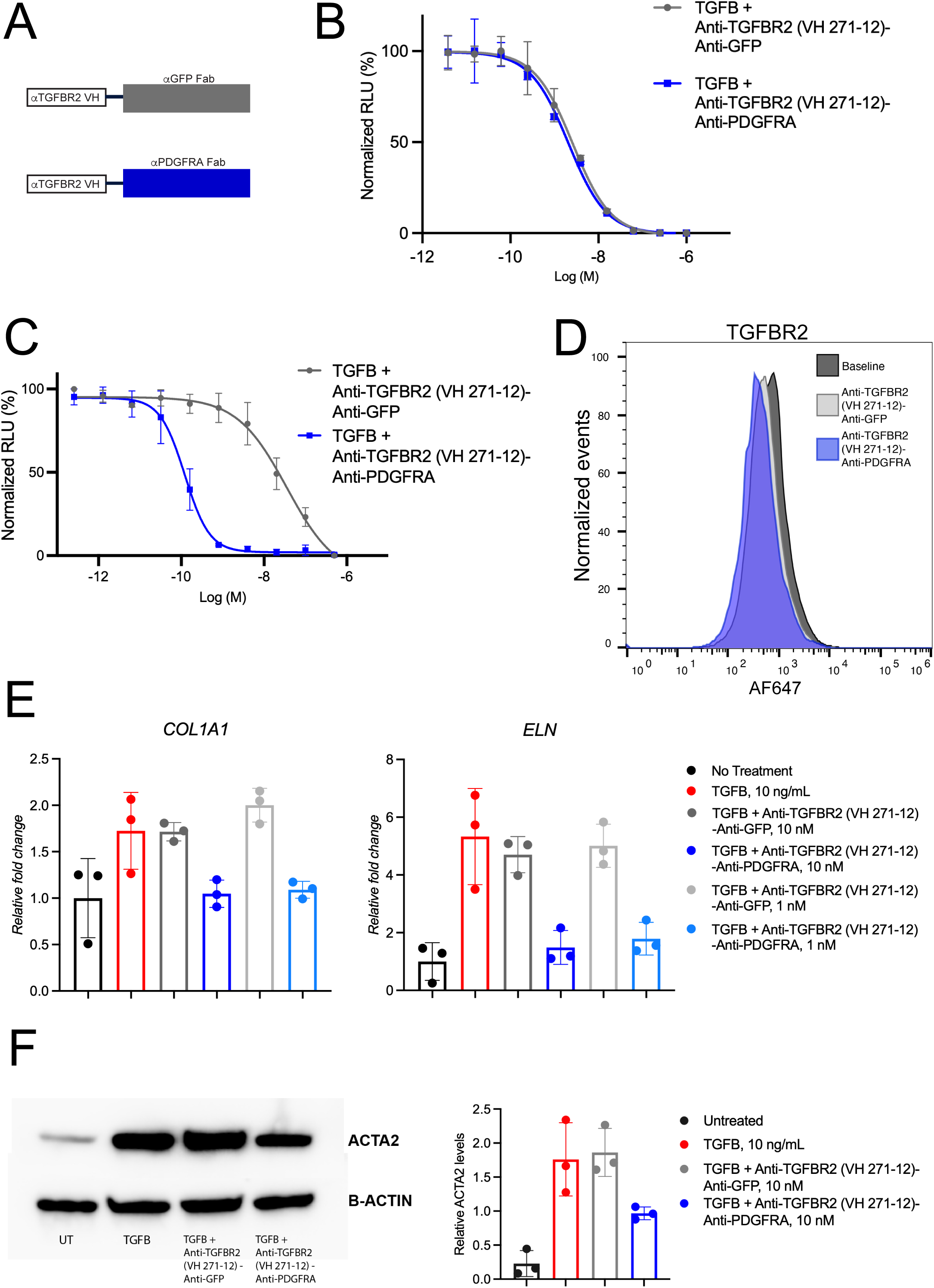
The PDGFRA-targeted TGFBR2 antagonist decreases signaling in human fibroblasts. (A) Schematic of the control and targeted TGFBR2 antagonists. (B, C) Activity of the untargeted and targeted antagonists in the SBE luciferase reporter assay in HEK293 cells (B) and human MRC5 fibroblasts (C). (D) Surface levels of TGFBR2 measured by flow cytometry after antagonist treatments in MRC5 fibroblasts. (E) RT-qPCR indicating ECM gene expression after antagonist treatments in MRC5 fibroblasts. (F) Western blotting for TGFB1-induced protein ACTA2 and the loading control B-ACTIN with quantification of the bands. All assays were completed in triplicate.

We assessed the activity of the targeted bispecific TGFBR2 antagonist in human embryonic pulmonary fibroblasts as well as HEK293 cells, which served as a negative control cell line without PDGFRA expression. In HEK293 cells, the targeted antagonist did not show greater activity than the untargeted control antagonist (Figure 3B). However, in the fibroblasts, the untargeted antagonist (VH271-12-aGFP) decreased TGFB1-induced signaling with an IC50 of 35 nM, and the PDGFRA targeted antagonist (VH271-12-aPDGFRA) was nearly 300-fold more potent, inhibiting TGFB1-induced activity with an IC50 of 120 pM (Figure 3C). Flow cytometry indicated that the untargeted antagonist alone modestly decreased surface levels of TGFBR2 whereas treatment with the anti-TGFBR2 VH + anti-PDGFRA Fab targeted antagonist further decreased surface TGFBR2 levels to isotype control levels (Figure 3D). In fibroblasts the targeted TGFBR2 antagonist inhibition of TGFβ signaling had functional effects on the expression of TGFB-responsive genes associated with ECM and fibrosis (Figure 3E, F). These results therefore provided an *in vitro* proof-of-concept for a cell-targeted TGFBR2 antagonist with potential for modulating fibrotic endpoints.

Even though TGM6-D3 and TGFB have overlapping binding epitopes on TGFBR2, TGM6-D3 had no inhibitory activity on its own in mouse fibroblasts within the tested concentration range (Fig. 1E). In contrast, a competitive human TGFBR2 antibody binder (VH271-12) with lower affinity to TGFBR2 than TGM6-D3 had modest antagonistic activity alone in human cells (Fig. 3C, VH271-12-aGFP curve). While this article was in preparation, a paper was published, in agreement with our findings, that TGM6 uses LRP1 as a targeting receptor leading to a LYTAC-mediated decrease in TGFBR2 levels (van Dinther et al., 2026). Furthermore, betaglycan (TGFBR3) was identified as an additional receptor for the TGM6-D3 domain that may inhibit its TGFBR2 binding (van Dinther et al., 2026). Whether this additional TGFBR3 binding might further suppress on-target off-tissue effects and increase cell selectivity, and whether other mechanisms might explain the heightened sensitivity of human cells to competitive TGFBR2 binders, warrants further consideration. It is also interesting to note that, although there was no enhancement of antagonist activity with the targeted molecule in the HEK293 cells, these cells were more sensitive to the untargeted ligand competitive antagonist than the fibroblasts (compare VH271-12-aGFP curves in Fig. 3B vs Fig. 3C). One possible explanation for this is that HEK293 cells express lower levels of TGFBR2. If this were to be the cause, it highlights an additional risk that some untargeted cells might be more sensitive to any molecule that contains a ligand competitive binder. Using a ligand non-competing TGFBR2 binder in the targeted antagonist may avoid this undesirable phenomenon.

Although the TGFβ pathway has been recognized as an attractive target for many diseases, on-target systemic toxicity has been a persistent roadblock to development of TGFβ pathway inhibitor therapeutics. As a strategy to increase the therapeutic window of TGFβ signaling antagonists, we generated a fully antibody-based TGFBR2 antagonist based on TGM6 that engages PDGFRA and has cell-type selective, enhanced antagonist activity in human fibroblasts. This work provides a framework for generating safer TGFβ signaling antagonists as therapies for cancer and fibrosis.

## Materials and Methods

### Molecular cloning

All proteins were expressed using the pcDNA3.1(+) mammalian expression vector (Thermo Fisher) with an N-terminal signal peptide. TGM6 was S16-T254 (NCBI MG429741) followed by a 6×His tag. TGM6-D3-Fc was S16-P102 (NCBI MG429741) followed by GGGGS and human IgG1-Fc containing L234A/L235A/P329G mutations (LALAPG). VH 271-12-TGM6-D4D5 was the VH binder 271-12 (clone DOM23h-271-12, WO2012/093125A1) followed by (G4S)×3 and TGM6-D4D5-6×His tag. VH 271-12-Fc was the VH 271-12 followed by GGGGS and human IgG1-Fc containing L234A/L235A/P329G mutations (LALAPG). VH 271-12-anti-GFP was the VH binder 271-12 followed by (G4S)×3 and the heavy chain Fab anti-GFP domains with a 6×His tag paired with the light chain Fab of the anti-GFP antibody. The anti-GFP binder was from an in-house screening campaign. VH 271-12-anti-PDGFRA was the VH binder 271-12 followed by (G4S) ×3 and the heavy chain of the anti-PDGFRA Fab binder (binder 2.1623.2, US2014/8697664) with a 6×His tag paired with the light chain Fab of the anti-PDGFRA Fab (2.1623.2, US2014/8697664). The amino acid sequence of human TGFBR2 used was N42-D159 (NCBI NP_003233.4). The protein was expressed as: human TGFBR2-GGGGS-Avi tag-GG-thrombin cleavage site-GGGGSGGGGS-human IgG1 Fc. The amino acid sequence of mouse TGFBR2 used was G42-D159 (NCBI NP_083851.3). The protein was expressed as: mouse TGFBR2-GGGGS-Avi tag-GG-thrombin cleavage site-GGGGSGGGGS-human IgG1 Fc.

### Protein production

All recombinant proteins were produced in Expi293F cells (Thermo Fisher Scientific) through transient transfection. The proteins were first affinity purified with MabSelect SuRe (Cytiva) or cOmplete His-tag purification resin (Sigma-Aldrich) and further purified with Superdex 200 Increase 10/300 GL (Cytiva) size-exclusion chromatography (SEC) with 1× HBS buffer (20 mM HEPES pH 7.4, 150 mM NaCl) except for recombinant human and mouse TGFBR2 which were polished with HiLoad 16/600 Superdex 200 pg (Cytiva) SEC with 2×HBS buffer (40 mM HEPES pH 7.4, 300 mM NaCl) on an AKTA pure system. The proteins were subsequently examined with SDS-polyacrylamide gel electrophoresis and estimated to have >90% purity.

### Binding assays

Binding kinetics were assessed by BLI biolayer interferometry with an OctetRed 96 instrument (Pall ForteBio, Fremont, CA). The monovalent KD was calculated according to a 1:1 fitting model. For TGFBR2 binding to TGM6, human TGFBR2-Fc or mouse TGFBR2-hFc diluted to 50 nM in running buffer (phosphate-buffered saline, 0.05% Tween-20, 0.5% bovine serum albumin, pH 7.2) was captured by Anti-hIgG Fc Capture (AHC) biosensors (Sartorius, Göttingen, Germany) and then dipped into a serial dilution of TGM6. Sensors were then dipped into running buffer to measure the dissociation of the interaction. To complete the binding competition assays, AHC sensors captured human TGFBR2-Fc or mouse TGFBR2-hFc, each diluted to 50 nM in running buffer. Sensors were then dipped into running buffer alone or TGFB1 (R&D Systems, 7754-BH-100/CF) diluted to 50 nM in running buffer, followed by TGM6 diluted to 50 nM in running buffer. To determine the affinity of VH 271-12 to human and mouse TGFBR2, VH 271-12 was diluted to 50 nM in running buffer and captured by the AHC sensor.

Sensors were then dipped into human or mouse TGFBR2 diluted to various concentrations in running buffer. Dissociation was measured by dipping the sensors into running buffer alone. For determination of competition between VH 271-12 and TGFB1, streptavidin (SA) biosensors (Sartorius, Göttingen, Germany) captured biotinylated recombinant humanTGFB1 (R&D Systems) diluted to 50 nM in running buffer. Sensors were then dipped into human TGFBR2-Fc diluted to 100 nM in running buffer or running buffer alone, followed by VH 271-12 diluted to 200 nM in running buffer.

### In vitro cell-based assays

Human TGFB1 (R&D Systems, 7754-BH-100/CF) was used at a concentration of 5 ng/mL in the luciferase and western blotting assays. TGFβ signaling activity was assessed using reporter cell lines expressing a luciferase gene under the control of a TGFB-responsive SMAD-binding element (SBE). NIH-3T3-SBE (Signosis, SL-0030), HEK293-SBE (BPS Biosciences, 60653), and MRC5-SBE cells (generated internally by transducing MRC-5 cells (ATCC, CCL-171) with a lentivirus carrying a SBE driven luciferase reporter transgene (Gentarget, LVP1714) and selected for puromycin resistance for 2-weeks) were maintained in DMEM supplemented with 10% fetal bovine serum (FBS), 1× HEPES, and 1× penicillin–streptomycin.

For activity assays, cells were seeded into the inner 60 wells of white 96-well plates at a density of 20,000–30,000 cells per well in maintenance medium. SBE reporter cells were plated on collagen-coated plates to enhance attachment. The following day, 50 µL of 2× antagonist solution was added to each well 30 minutes before stimulation with recombinant human TGFB1. Cells were incubated for 16–20 h at 37 °C. After treatment, the culture medium was removed, and the cells were immediately lysed in 100 µL of 1× lysis buffer (Promega, E153A) with shaking at room temperature for 10 minutes. Lysates (20 µL) were transferred to white opaque 96-well plates containing 10 µL Luciferin Detection Reagent (Promega, E1501). Luminescence was measured with an EnVision plate reader according to the manufacturer’s instructions.

For SMAD2/3 and phospho-SMAD2/3 western blotting on NIH-3T3 cells, where indicated, TGM6 was applied 30-minutes before treatment with TGFB1, and molecule treatments proceeded overnight (∼16-18 hours) at 37 °C. Cells were dissociated and lysed with Pierce RIPA buffer (Thermo Scientific, 89901) containing protease and phosphatase inhibitors (Thermo Scientific, 1861281) before boiling in 1× Laemmli buffer followed by SDS-PAGE. Proteins were transferred to nitrocellulose membranes and blotted for SMAD2/3 (Cell Signaling Technologies, #8685) and bis-phospho-SMAD2/3 (Cell Signaling Technologies, #8828), then visualized with an HRP-conjugated secondary antibody (Abcam, AB205718) via chemiluminescence with SuperSignal West Pico PLUS Chemiluminescent Substrate (Thermo Scientific, 34578). For ACTA2 western blotting, equal numbers of cells were directly lysed in 1× Laemmli buffer followed by SDS-PAGE. Proteins were transferred to nitrocellulose membrantes and blotted for ACTA2 (Abcam, AB7817) and B-ACTIN (Abcam AB8226) and visualized as described above.

### RT-qPCR on MRC5 fibroblasts

Following overnight (∼18-hours) treatment incubations on MRC5 fibroblasts at 37 °C, cells were dissociated, and RNA was isolated with the Zymo Research Direct-Zol RNA microprep kit (R2660), and the Invitrogen Superscript IV VILO master mix (Invitrogen, 11756050) cDNA synthesis kit was used for cDNA synthesis. TaqMan Fast Advanced Master mix (Thermo Scientific 444554) was used for qPCR. The ddCt method was used for relative quantification with respect to the housekeeping gene, *GAPDH*. The following TaqMan probes were used: GAPDH, Hs02786624_g1; COL1A1, Hs00164004_m1; ELN, Hs00355783_m1.

### Flow Cytometry

Cells were dissociated and washed in complete medium, then resuspended in cold flow buffer (2% FBS in PBS) and kept on ice. After Fc receptor blocking (mouse FcR block, Miltenyi Biotec 130-092-575; human Fc receptor block, 130-059-901), cells were incubated with primary antibodies to TGFBR2 (R&D Systems goat anti-mouse TGFBR2 (AF532) or goat anti-human TGFBR2 (AF241)) or a goat IgG isotype control antibody (R&D Systems, AB-108-C) followed by washes with flow buffer and incubation with a fluorescently-labeled secondary antibody followed by washes in flow buffer. Cells were analyzed on a Sony SH800 cell sorter and live-dead gated with DAPI. Data are presented as normalized events with the indicated secondary antibody fluorescence signal. In all molecule treatment experiments assessed by flow cytometry, cells were treated with the indicated molecules for 30-minutes, unless specified otherwise, at 37 °C, then washed with PBS, dissociated with Trypsin + EDTA, and quenched with cold complete medium and immediately placed on ice, washed with cold flow buffer, and prepared as described above for analysis. The only exception to this was with the washout experiments, in which TGM6 treatment incubation was for 30 minutes followed by medium replacement absent TGM6 for 24-48 hours. The dynamin inhibitor Dynasore was obtained from Selleckchem (S8047). Dynasore was applied for 30 minutes before TGM6 treatment application and was maintained in the medium throughout the duration of the experiment.

### Quantification

Cell based in vitro assays (reporter assays, western blots, RT-qPCR) were completed in a minimum of biological triplicates except the VH 271-12-TGM6-D4D5 fusion protein studies in MRC5 human fibroblasts which were done in duplicate.

## Acknowledgments

We thank our scientific founders, Christopher Garcia and Michael Elowitz, and our Scientific Advisory Board members Dean Sheppard and Andy Hinck for their guidance and support. We thank Craig Parker for helpful discussions and critical reading of this manuscript. This work was funded by TCGFB, Inc.

